# CTRP7 as a Molecular Biomarker for Predicting Responsiveness to Pulmonary Vasodilators: Insights from Human and Animal Studies in Pulmonary Arterial Hypertension

**DOI:** 10.1101/2023.11.05.565725

**Authors:** Kaito Yamada, Taijyu Satoh, Nobuhiro Yaoita, Saori Yamamoto, Haruka Sato, Yusuke Yamada, Kohei Komaru, Naoki Chiba, Takashi Nakata, Kotaro Nochioka, Hisashi Oishi, Satoshi Miyata, Yoshinori Okada, Satoshi Yasuda

## Abstract

**Background:** Pulmonary arterial hypertension (PAH) is a life-threatening condition. Although pulmonary vasodilators have shown promise in managing PAH, the overall prognosis improvement is modest, partly because of the absence of a biomarker for guiding their selection. This study aimed to identify the molecular-based predictors of responsiveness to pulmonary vasodilators by combining human and basic studies.

**Methods and Results:** RNA sequencing was conducted on cultured pulmonary artery smooth muscle cells (PASMCs) from patients with and without PAH, identifying variations in 2,076 genes. The candidates were narrowed down to established clinical biomarkers, and their plasma levels were assessed in PAH (n = 123) and non-PAH patients (n = 25). C1q/TNF-related protein 7 (CTRP7) exhibited elevated expression in PASMCs (5.37 Log2 fold change, p < 0.01), and the plasma of patients with PAH compared to non-PAH (28.2 (9.3– 101.2) vs. 9.32 (4.6–18.6) ng/ml, p < 0.01). A plasma assessment revealed a significant correlation between CTRP7 and interleukin 6 (IL-6) levels (r = 0.544, p < 0.001). Chromatin immunoprecipitation demonstrated that IL-6 upregulated CTRP7 expression in PASMCs. Among pivotal factors, CTRP7 reduced prostacyclin analog receptor (PTGIR) expression through Rab5a-mediated internalization, resulting in diminished responsiveness to selexipag (a prostacyclin analog). Incorporating the human study, hypoxic PH mice demonstrated reduced PTGIR expression in the pulmonary arteries, which correlated with limited responses to selexipag treatment (low cardiac output and persistent pulmonary artery resistance), which was mitigated by silencing CTRP7 expression in the pulmonary arteries using adeno-associated virus 6.

**Conclusions:** Among the candidates with increased expression in patients with PAH based on RNA sequencing of PASMCs, CTRP7 demonstrated elevated plasma levels compared to non-PAH. CTRP7 regulated PTGIR internalization via inflammation and Rab5a, influencing responsiveness to selexipag. CTRP7 emerged as a compelling biomarker for predicting responsiveness to prostacyclin analogs, potentially advancing the development of innovative PAH treatment strategies.

**Novel and Significance:** *What Is Known?:*

- Pulmonary arterial hypertension (PAH) is a life-threatening vascular disease with limited treatment options.
- Although pulmonary vasodilators have shown promise in managing PAH, the overall improvement in prognosis is modest.
- Further research is needed to understand the molecular features of pulmonary vasodilators and identify biomarkers for appropriate selection.

*What New Information Does This Article Contribute?:*

- C1q/TNF-related protein 7 (CTRP7) was found to be elevated in the pulmonary artery smooth muscle cells (PASMCs) and plasma of patients with pulmonary arterial hypertension (PAH).
- This study established a connection between inflammation and pulmonary vascular responsiveness via CTRP7-mediated internalization and degradation of the prostacyclin analog receptor.
- This study indicated the possibility of using plasma CTRP levels as a clinical biomarker to monitor pulmonary responsiveness, which can contribute to the development of novel treatment strategies for patients with PAH. Pulmonary arterial hypertension (PAH) is a life-threatening and enigmatic lung vascular disorder characterized by progressive pulmonary vascular remodeling and vasoconstriction. Despite significant advancements in the treatment of PAH using pulmonary vasodilators, some individuals display limited responses, primarily due to a lack of understanding of the molecular features of each pulmonary vasodilator and the absence of a biomarker for appropriate selection. The present study aimed to identify promising biomarkers for predicting a patient’s responsiveness to pulmonary vasodilators through a two-step approach, involving RNA sequencing of PASMCs and the measurement of plasma levels. CTRP7 emerged as a prominent biomarker, displaying increased expression in both PASMCs and plasma obtained from patients with PAH. Mechanistically, IL-6 transcriptionally modulated CTPR7, leading to the internalization and degradation of PTGIR, resulting in a reduced response to selexipag (prostacyclin analog) in PASMCs. Integrating the human studies, a hypoxic mouse model exhibited lower PTGIR expression in pulmonary arteries, correlating with limited responses to selexipag treatment, which were ameliorated by silencing CTRP7 expression in pulmonary arteries using adeno-associated virus 6 transduction. This study collectively demonstrates that CTRP7 emerges as a compelling biomarker for predicting responsiveness to prostacyclin analogs, with the potential to advance the development of innovative PAH treatment strategies.

## Introduction

Pulmonary arterial hypertension (PAH) is a life-threatening and enigmatic lung vascular disorder characterized by progressive pulmonary vascular remodeling and vasoconstriction, which ultimately results in right ventricular failure and mortality.^1–3^ Presently, management of PAH primarily involves the use of pulmonary vasodilators that target the endothelin, nitric oxide (NO)-soluble guanylate cyclase (sGC)-cyclic guanosine monophosphate (cGMP) signaling, or the prostacyclin pathways. Both monotherapy and combination regimens involving these pulmonary vasodilators have significantly enhanced the clinical outcomes of patients with PAH.^4–8^

Despite significant advancements in the treatment of PAH, mortality rates among patients with PAH remain distressingly high, with some patients displaying limited responses to vasodilator therapies,^9, 10^ and a substantial proportion, approximately 30–40%, of patients with PAH, continuing to exhibit mean pulmonary artery pressures exceeding 45 mmHg.^9^ Furthermore, the use of vasodilators is associated with an elevated incidence of adverse events, which can limit their long-term tolerability. Indeed, a substantial proportion of patients undergoing oral or inhaled monotherapy continue to experience severe pulmonary hypertension (PH), characterized by a mean pulmonary artery pressure exceeding 45 mmHg.^9^ This is plausibly due to the lack of adequate evidence for the appropriate application of vasodilators, and understanding of the molecular features of each vasodilator. The European society of cardiology / European respiratory society (ESC/ERS) guideline provides the algorism for selection of pulmonary vasodilators following risk classification, including WHO functional class, 6 minutes walking distance, and plasma level of brain natriuretic peptide. Patients with high-risk or residual PAH, even after treatment with Phosphodiesterase-5 inhibitors or endothelin receptor antagonists, should be administered prostacyclin receptor analogs, such as selexipag, 6reprostinil, or epoprostenol. In clinical practice, the treatment strategy relies on the individual experiences of each physician, which leads to empiric-multiple-use of pulmonary vasodilators, resulting in insufficient clinical course and unresolved issues in patients with PAH.

Despite the substantial body of evidence for abnormal pulmonary arterial remodeling in PAH, the mechanisms underlying the responsiveness to vasodilators remain incompletely elucidated in basic research. Recently, therapies such as Riociguat or Vericiguat, which target the NO-sGC axis, have garnered attention in the context of heart failure and PH. Explorations into the molecular mechanisms involved have revealed insights into factors such as reduced NO production in diabetes,^10^ and the oxidation of sGC.^11^ Prostacyclin agonists such as selexipag, treprostinil, and epoprostenol downregulate prostacyclin analog receptor expression in the pulmonary arteries of PAH patients.^12^ However, the molecular evidence is not sufficient, and practical tests or biomarkers capable of evaluating the responsiveness to vasodilator therapies are yet to be identified.

Therefore, in this study, we aimed to elucidate the mechanisms underlying responsiveness to pulmonary vasodilators by conducting multiple analyses utilizing pulmonary artery smooth muscle cells (PASMCs) and plasma samples obtained from patients diagnosed with PAH. The C1q/TNF-related proteins (CTRPs) are a family of secreted proteins produced by various organs that share functional similarities with adiponectin. Dysregulation of CTRPs has been linked to metabolic abnormalities such as obesity, type 2 diabetes, and cardiovascular disease.^13^ Among the CTRPs, CTRP7 is a well-established plasma biomarker for indicating insulin resistance.^14^ Despite its high expression in the lungs, the role of CTRP7 in the pathogenesis of PAH has remained unknown.^15^

By combining data of both human and basic studies, we present evidence demonstrating that CTRP7 exerts a negative regulatory effect on PTGIR expression via Rab5a, and consequently, impairs the response to selexipag in cultured PASMCs and hypoxic mice.

## Methods

### Clinical Data

We conducted a retrospective investigation involving 98 patients with precapillary pulmonary hypertension who had undergone right heart catheterization between January 2015 and December 2023. The diagnosis was established through a combination of echocardiography, computed tomography, spirometry, ventilation/perfusion lung scans, and right heart catheter examination, adhering to the guidelines set forth by the European Society of Cardiology and European Respiratory Society in both 2015 and 2022.^4, 16^ The diagnostic criteria included a resting mean pulmonary artery pressure (mPAP) measured via right cardiac catheterization of ≥20 mmHg and a pulmonary artery wedge pressure (PAWP) of ≤15 mmHg. Patients who underwent catheterization but did not exhibit pulmonary hypertension were included as the non-PAH control group (n=25). Exclusions from the study comprised patients with significant valvular heart disease (defined as moderate-to-severe stenosis or moderate-to-severe regurgitation) and those who had undergone heart or lung transplants.

All protocols using human specimens were approved by the Institutional Review Board of Tohoku University, Sendai Japan. The protocol accords with the ethical guidelines of Declaration of Helsinki. This study was approved by the Medical Ethics Review Committee (approval no., 2021-1-208) which waived the requirement for informed consent because of the retrospective nature of the study. We also applied Opt-out method to obtain consent on this study.

### Animal Experiments

All animal experiments were conducted in compliance with the protocols approved by the Tohoku University Animal Care and Use Committee (Approval No. 2021-ido-073) following the ARRIVE guidelines. Hypoxia-induced PH models were employed to evaluate PH development in mice (19). Eight-week-old male wild-type (WT) mice, fed a normal chow diet, were subjected to either hypoxia (10% O2) or normoxia for a duration of 4 weeks, as previously described.^17^ In brief, mice exposed to hypoxia were housed in an acrylic chamber supplied with a non-recirculating gas mixture of 10% O_2_ and 90% N_2_, utilizing an adsorption-type oxygen concentrator for exhaust air (Teijin, Tokyo, Japan). Concurrently, normoxic mice were housed in room air (21% O2) under a 12-hour light-dark cycle. After 2 weeks of hypoxic or normoxic exposure, mice were administered with either 1 mg/kg of Selexipag or a vehicle containing 1% dimethyl sulfoxide (DMSO) daily.^18^ Following 4 weeks of hypoxic or normoxic exposure, mice were anesthetized using 1.0% isoflurane. To evaluate the development of pulmonary hypertension, measurements were taken for right ventricular systolic pressure (RVSP), right ventricular hypertrophy (RVH), and pulmonary vascular remodeling. For right heart catheterization, a 1.2-F pressure catheter (SciSense Inc., Ontario, Canada) was inserted into the right jugular vein and advanced into the right ventricle to measure RVSP. All data were recorded and analyzed using the Power Lab data acquisition system (AD Instruments, Bella Vista, Australia) and were averaged over 10 sequential beats. In the Sugen/hypoxia model, mice and rats (male Sprague-Dawley, aged 7-10 weeks) were subcutaneously injected with the VEGF-receptor inhibitor SU5416 (Sigma-Aldrich) at a dosage of 20 mg/kg body weight under isoflurane anesthesia. Subsequently, they were exposed to hypoxia (10% O2) for a duration of 3 weeks, following established procedures.

### Statistical Analyses

The results are presented as the mean ± standard deviation (SD). To compare means between two groups with equal variances, a two-tailed Student’s t-test was employed. Alternatively, the Mann–Whitney U test was utilized for comparisons between two groups with unequal variances. To evaluate the mean responses associated with the two main effects of distinct groups, a two-way analysis of variance (ANOVA) was conducted, followed by Tukey’s HSD (honestly significant difference) test for multiple comparisons. The relationship between plasma CTRP7 levels and IL-6 was analysed using Pearson’s rank correlation coefficient. Statistical significance was assessed with JMP 12 (SAS Institute Inc., Cary, USA) or GraphPad Prism 7 (GraphPad Software, San Diego, USA). All reported p-values are two-tailed, with a p-value of less than 0.05 considered statistically significant.^19^

## Results

### Upregulation of CTRP7 in Pulmonary Artery Smooth Muscle Cells in Pulmonary Arterial Hypertension

RNA sequencing was performed on PASMCs obtained from both patients with and without PAH, and variations in 2, 076 genes were identified (**Figure 1A**). Considering the emphasis on the clinical significance and identification of a potential biomarker for PAH, the prospective biomarker candidate proteins encoded by these genes were narrowed down to those with an established biomarker status in existing clinical research (**Figure 1A** and **1B**). Among these candidates, CTRP7 demonstrated a significant increase in mRNA expression ­­­in PAH PASMCs compared to that in non-PAH PASMCs (Log2 fold change 5.37) (**Figure 1A**). Furthermore, CTRP7 was highly expressed in the plasma of patients with PAH, surpassing that of candidates such as PLA2G2A or TNC (**Figure 1C** and **Supplemental Figure IA**). To confirm the expression site of CTPR7 in vasculatures, the expression data of ACTA2 (representing smooth muscle), and CTPR7 gene was extracted from Tabula Muris project ^20^, which investigated various murine tissues using single-cell RNA seq experiments (**Figure 1D**). The graph suggested that CTPR7 expresses in stromal cells rather than endothelial cells or lymphocyte in the single-cell analysis of mouse lung (**Figure 1D**). The cluster expressing ACTA2 indicates the smooth muscle cells, which was overlapped with the area of CTRP7 expresses, suggesting CTPR7 expresses from smooth muscle cells. Consistent with the mRNA analyses, CTRP7 protein levels were notably elevated in PASMCs derived from patients with PAH (**Figure 1E**), as well as in lung homogenates harvested from animal models of PH, including the hypoxic mouse model and the Sugen/hypoxia rat model (**Figure 1F** and **G**), when compared with their respective controls. CTRP7 expression was not altered in the adipose tissue, liver, left and right ventricle of hypoxic mice compared to those of normoxic mice (**Supplemental Figure IB**). In line with these findings, CTRP7 plasma concentrations in patients with PAH (n = 98) were compared with those in patients without PH or cardiovascular disease (**Figure 1E**). The baseline characteristics of the two groups, including age, sex, and comorbidities, were comparable (**Supplemental Table I**). CTRP7 was significantly higher in the plasma of patients with PAH compared with those without [PAH vs non-PAH: 9.3 (4.6–18.6) vs 26.7 (9.3–101), p = 0.0001] (**Figure 1E**). Furthermore, CTRP7 levels showed a relatively higher trend in patients with connective tissue disease-associated PAH (CTD-PAH) compared with patients with non-CTD-PAH [CTD-PAH vs. non-CTD-PAH: 33.5 (14.8–109.1) vs. 18.2 (8.4–45.2) ng/ml, p = 0.07], although this difference did not reach statistical significance, which is noteworthy considering the well-established association of connective tissue diseases with inflammation.

**Figure 1.**
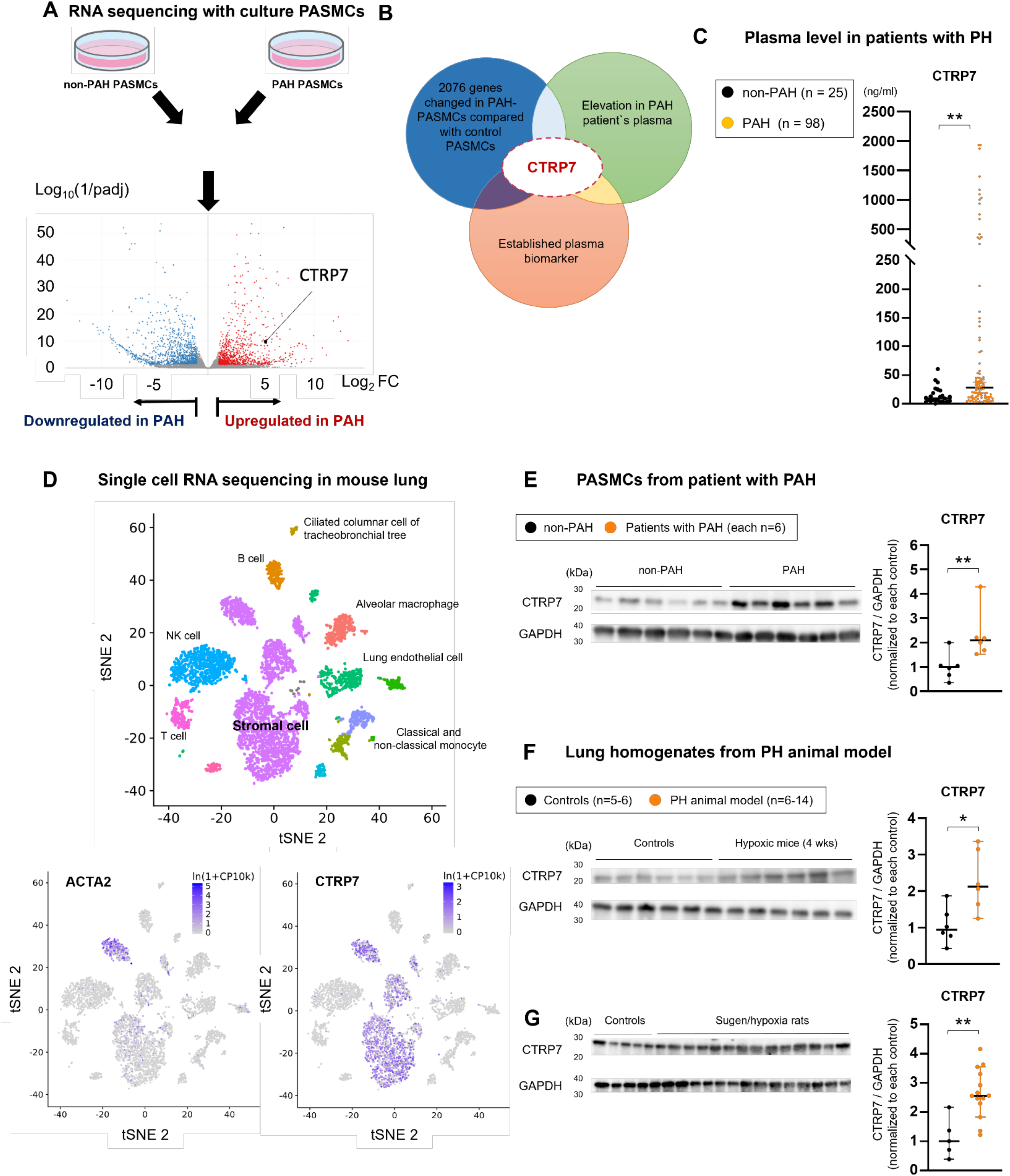
Upregulation of C1q/TNF-related protein 7 in Patients with PAH and in PH Animal Models. **(A)** RNA sequencing of PASMCs from patients with PAH and those from non-PAH individuals. **(B)** Selection of CTRP7 as a candidate gene based on RNA sequencing and comprehensive plasma assessment. **(C)** Elevated CTRP7 plasma levels in patients with PAH (n = 98) compared with that in non-PAH individuals (n = 25). **(D)** Expression of ACTA2 (actin alpha 2) and CTRP7 in murine tissues. The data was obtained from Tabula Muris project. Expression in corresponding tissue origin of all cells (upper graph). The cells were clustered using t-SNE (t-distributed Stochastic Neighbor Embedding) algorithm based on their transcriptome. ACTA2 is a key smooth muscle contractile protein, and its expression in the graph indicates the smooth muscle clustering in stromal cells. CTRP7 expresses in almost all stromal cells including smooth muscle cells. **(E)** Representative western blot and quantification of CTRP7 in PASMCs from patients with PAH. **(F)** Representative western blot and quantification of CTRP7 in lung homogenates from hypoxic mice, **(G)** Sugen/hypoxia rats, and their corresponding controls. PASMCs, pulmonary artery smooth muscle cells; PAH, pulmonary arterial hypertension; CTRP7, C1q/TNF-related protein The results are presented as mean ± SEM. Statistical comparisons of the parameters were conducted using a 2-tailed Student’s t-test or Welch’s t-test, as appropriate. The significance levels are denoted as follows: *p < 0.05, **p < 0.01, and ***p < 0.001.

### CTRP7 Downregulates Prostaglandin I2 receptor in PASMCs of patients with PAH and in a Rodent Animal Model

To evaluate the role of CTRP7, we selectively downregulated it in PASMCs utilizing siRNA. CTRP7 downregulation did not alter levels of cell proliferation factors such as Cycline D1(CCND1) and proliferating cell nuclear antigen (PCNA) (**Supplemental Figure IIA**). To explore the role of CTRP7 in the vasoreactivity of the pulmonary arteries, we evaluated the key targeting factors in the three major mechanisms of pulmonary vasodilators: the prostaglandin (PG)I2, NO-sGC-cGMP axis, and endothelin pathways. CTRP7 downregulation led to a significant increase in PTGIR expression of PASMCs, while the expression of sGC β subunit and endothelin receptor type A were not significantly altered (**Figure 2A**). Similarly, PAH PASMCs exhibited higher CTRP7 expression and lower PTGIR expression than non-PAH PASMCs (**Figure 2B**), which were recovered by CTRP7 downregulation (**Figure 2C**). These findings indicate that CTRP7 is upregulated in PAH PASMCs and regulates PTGIR expression. Immunofluorescence analyses showed that PTGIR was significantly decreased in the pulmonary arteries of patients with PAH compared to that in those of patients without PAH (**Figure 2D**). Similarly, PTGIR expression was significantly lower in the pulmonary arteries of hypoxic mice and Sugen/hypoxia rats than in the corresponding controls **(Figure 2E** and **2F**).

**Figure 2.**
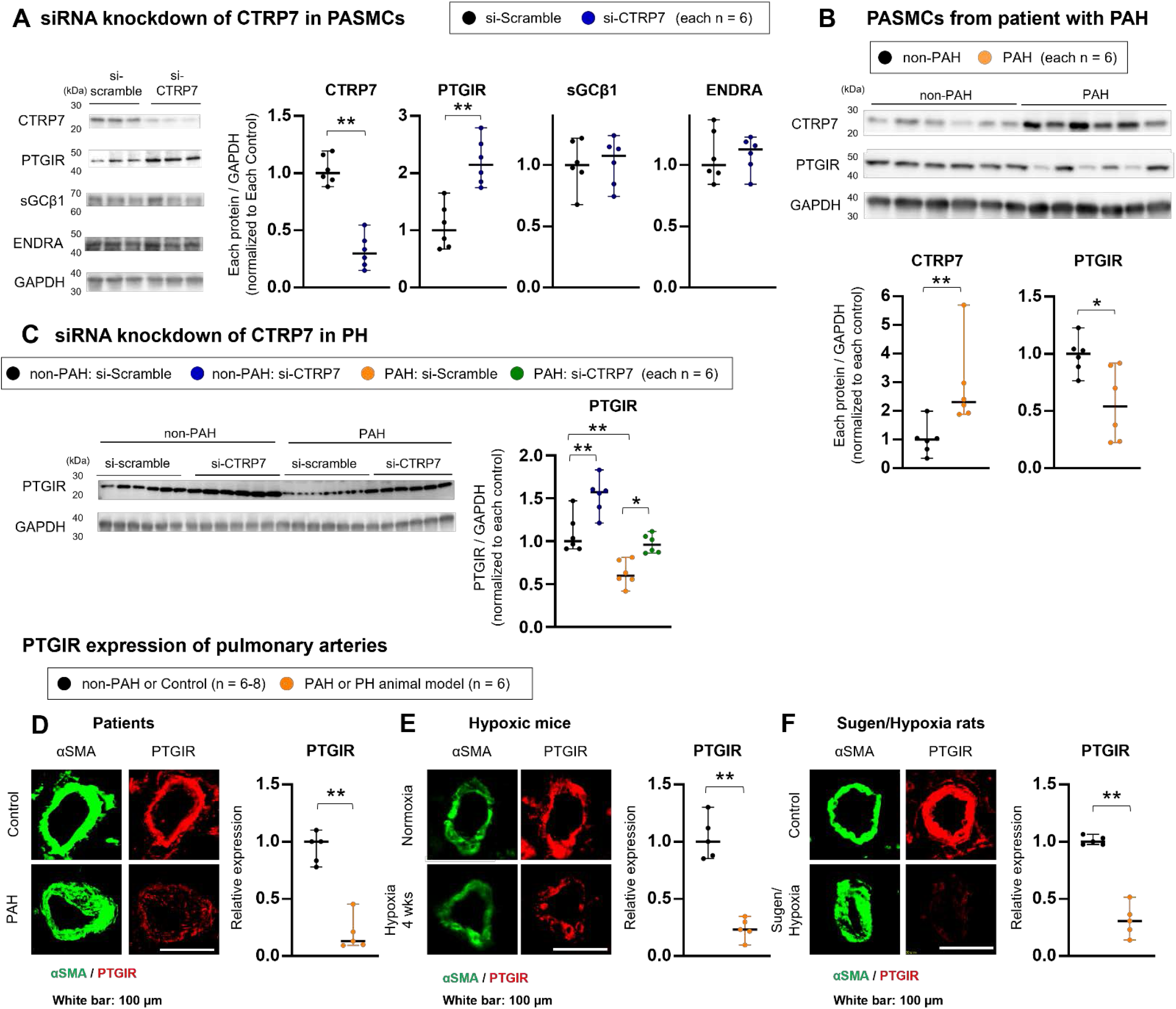
CTRP7 Downregulates Prostacyclin Analog Receptor in Patients with PAH and in Rodent Animal Models. **(A)** Representative western blot and quantification of PTGIR, sGCβ1, ENDRA, and GAPDH in human PASMCs transfected with si-CTRP7 or si-Scramble. **(B)** Representative western blot and quantification of CTRP7, PTGIR, and GAPDH in PASMCs from patients with PAH and from non-PAH individuals. **(C)** Representative western blots and quantification of PTGIR in PASMCs from patients with PAH and from non-PAH individuals transfected with si-CTRP7 or si-Scramble. **(D-F)** Representative images of immunostaining for αSMA (green) and PTGIR (red) in the distal PAs of patients with PAH, hypoxic mice (10% O_2_ for 4 weeks), Sugen/hypoxia rats, and corresponding controls. Scale bars = 100 μm. PTGIR, prostacyclin analog receptor; sGCβ1, soluble guanylate cyclase β1 subunit; ENDRA, endothelin receptor type A; PASMCs, pulmonary artery smooth muscle cells; CTRP7, C1q/TNF-related protein 7; PAH, pulmonary arterial hypertension; PAs, pulmonary arteries Data are presented as mean ± SEM. Statistical comparisons were conducted using the 2-tailed Student’s t-test, Welch’s t-test, or 2-way ANOVA, followed by Tukey’s honestly significant difference test for multiple comparisons. Significance levels are represented as *p < 0.05, **p < 0.01, and ***p < 0.001.

### Interleukin 6 transcriptionally Regulates CTRP7 Expression in PASMCs

Considering the well-established role of inflammation in the pathogenesis of PAH,^21^ we postulated that inflammation may play a crucial role in the elevation of CTRP7 expression observed in patients with PAH. Comprehensive plasma analyses revealed relative increases in interleukin (IL)-6, IL-8, IL-10, and TNFα in the plasma of both patients with non-CTD PAH and those with CTD-PAH (**Figure 3A** and **Supplemental Figure IIIA**). Notably, patients with CTD-PAH exhibited relatively higher plasma levels of IL-6 and CTRP7 than those with non-CTD-PAH (**Figure 3A**). Among these cytokines, IL-6 levels were significantly correlated with CTRP7 levels (r = 0.544, p = 0.0001) (**Figure 3B** and **Supplemental Figure IIIB**).

**Figure 3.**
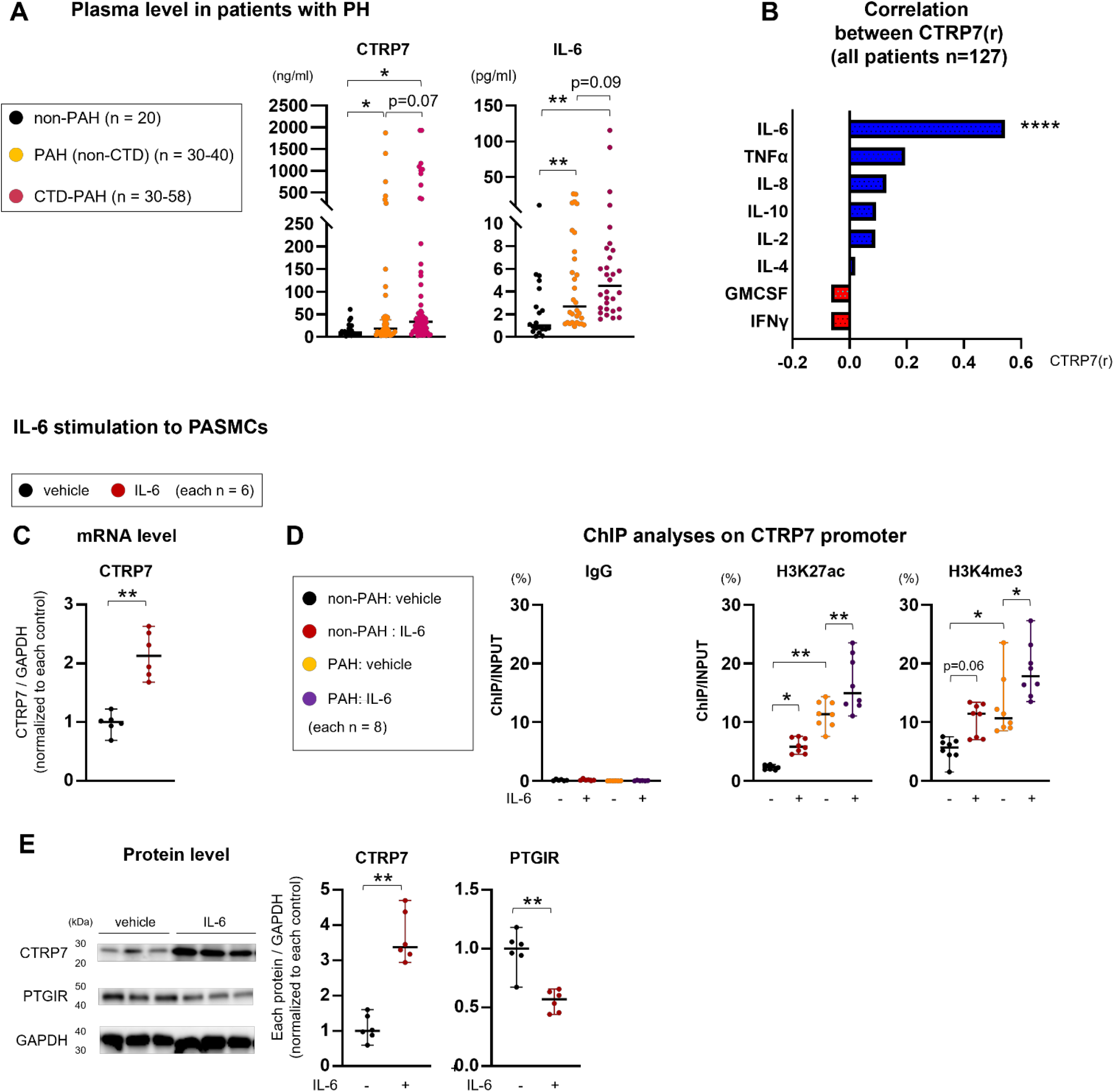
IL-6 Increases CTRP7 Expression via Transcriptional Mechanisms in PASMCs. **(A)** Plasma levels of CTRP7 and IL-6 in patients with non-CTD-PAH (n = 30–40), those with CTD-PAH (n = 30–58), and in non-PAH individuals (n = 20, 20). **(B)** Correlation between CTRP7 and plasma levels of inflammatory factors in patients with PAH and non-PAH individuals (n =123). **(C)** mRNA expression of CTRP7 in human PASMCs treated with IL-6 (20 ng/ml) or vehicle for 24 h. **(D)** Chromatin immunoprecipitation targeting H3K27 acetylation and H3K4 methylation of CTRP7 promoter in PASMCs treated with IL-6 (n = 8). **(E)** protein expression of CTPR 7 and PTGIR in human PASMCs treated with IL-6 (20ng/ml) for 24 hours (n = 6). CTRP7, C1q/TNF-related protein 7; IL-6, Interleukin-6; non-CTD-PAH: non-connective tissue disease-associated pulmonary arterial hypertension; PASMCs, pulmonary artery smooth muscle cells; H3K27, histones 3 lysine 27; H3K4, histone 3 lysine 4 Data are presented as mean ± SEM. Statistical comparisons were conducted using the 2-tailed Student’s t-test, Welch’s t-test, 1-way ANOVA, or repeated-measures 2-way ANOVA, followed by Tukey’s honestly significant difference test for multiple comparisons. Significance levels are indicated as *p < 0.05, **p < 0.01, and ***p < 0.001.

To elucidate the mechanistic role of IL-6 in the regulation of CTRP7 expression, we co-cultured PASMCs with recombinant IL-6 (20 ng/ml) for 24 hours. IL-6 stimulation significantly increased CTRP7 mRNA expression (**Figure 3C**). Chromatin immunoprecipitation assay was performed to investigate the transcriptional regulation of CTRP7 by IL-6. We predicted the promoter regions to be located within the first 1000 bp of the CTRP7 transcriptional starting site as previously described (**Supplemental Table II**).^22^ PASMCs treated with IL-6 from both non-PAH individuals and from patients with PAH exhibited higher levels of histone 3 lysine 27 (H3K27) acetylation and histone 3 lysine 4 (H3K4) methylation in the CTRP7 promoter region (**Figure 3D**).

Western blotting revealed that IL-6 stimulation significantly increased CTRP7 expression and reduced PTGIR expression (**Figure 4A**). Furthermore, the IL-6-induced decrease in PTGIR expression was attenuated by siRNA-mediated downregulation of CTRP7 (**Figure 4A**). In a hypoxic mouse model of PH, levels of CTRP7 substantially increased in both the plasma and lung tissue at the onset of hypoxic exposure or following monocrotaline stimulation, along with an increase in IL-6 and a decrease in PTGIR expression in the pulmonary arteries (**Figure 4B, 4C,** and **4D**). During the chronic phase of hypoxic conditions, these reactions relatively decreased but were sustained at a high level. A similar pattern was observed for monocrotaline-induced PH-model rats (**Supplemental Figure IVA**). Collectively, these findings indicated that CTRP7 regulates PTGIR expression via IL-6.

**Figure 4.**
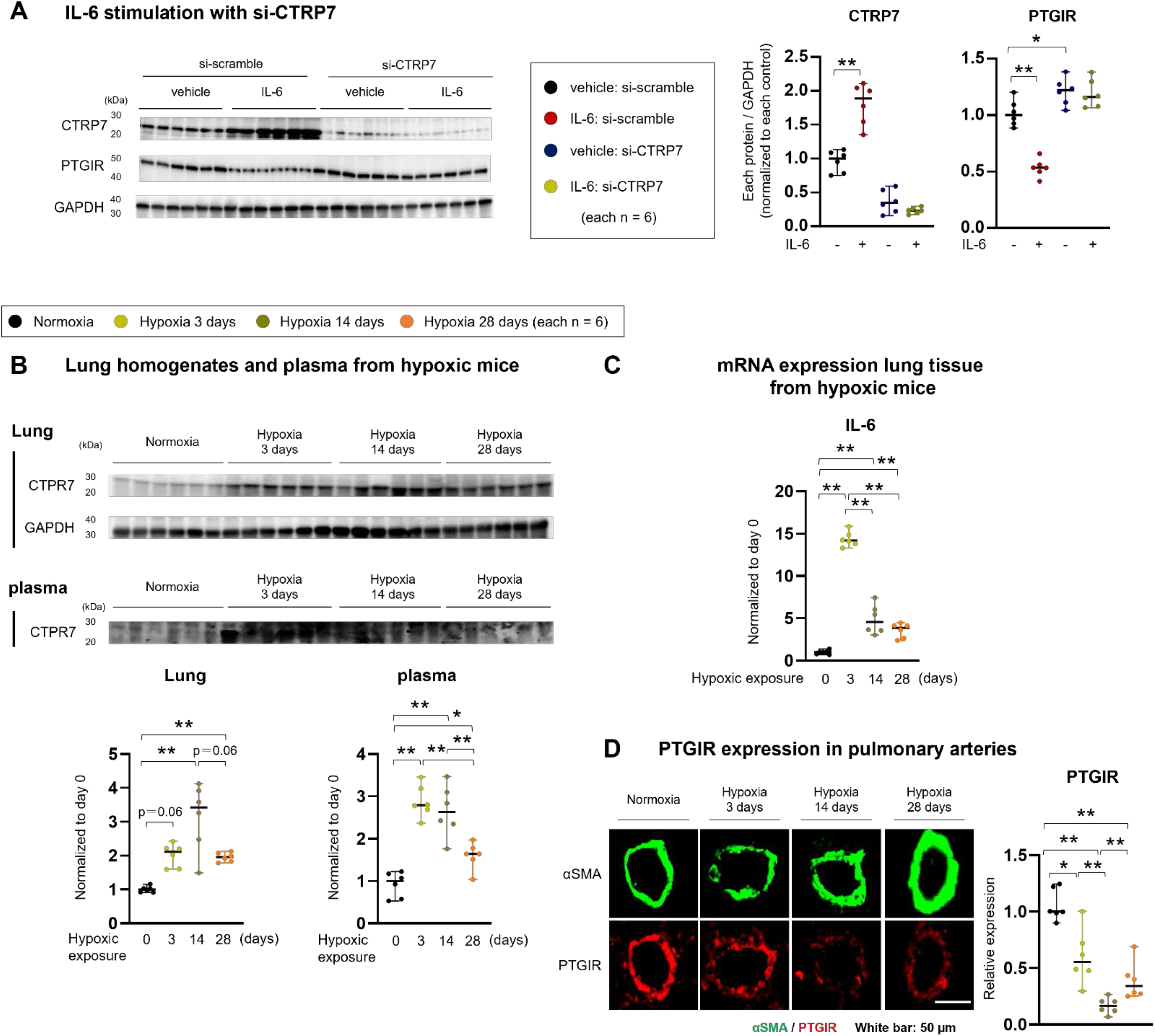
IL-6 Regulates Expression of CTRP 7 and Prostacyclin Analog Receptor in PASMCs and Lungs of PH Animal Models. **(A)** Representative western blot and quantification of CTRP7 and PTGIR in PASMCs with transfection of si-CTRP7 or si-Scramble, treated with IL-6 (20 ng/ml) or vehicle. **(B)** Representative western blot and quantification of CTRP7 and GAPDH in lung homogenates or plasma harvested from hypoxic mice (n = 6). **(C)** mRNA expression in the lungs of hypoxic mice (n = 6). **(D)** Representative immunofluorescent images and quantification of αSMA and PTGIR in pulmonary arteries of hypoxic mice (n = 6) CTRP7, C1q/TNF-related protein 7; PTGIR, prostacyclin analog receptor; PASMCs, pulmonary artery smooth muscle cells; IL-6, Interleukin-6; αSMA, alpha smooth muscle actin. Data are presented as mean ± SEM. Statistical comparisons were conducted using the 2-tailed Student’s t-test, Welch’s t-test, 1-way or 2-way ANOVA, followed by Tukey’s honestly significant difference test for multiple comparisons. Significance levels are indicated as *p < 0.05, **p < 0.01, and ***p < 0.001.

### Inflammation Induces Internalization of Prostacyclin Analog Receptor with Rab5a in PASMCs

Rab5a is known to control the vesicle-mediated transport of plasma membrane proteins to the endosomal compartment.^23^ We observed an increase in Rab5a expression in PAH PASMCs, which was significantly attenuated by siRNA-mediated downregulation of CTRP7 (**Figure 5A** and **B**). Consistent with these findings, co-incubation with recombinant CTRP7 significantly decreased PTGIR expression in non-PAH PASMCs (**Figure 5C**), pointing to the role of CTRP7 in regulating Rab5a expression.

**Figure 5.**
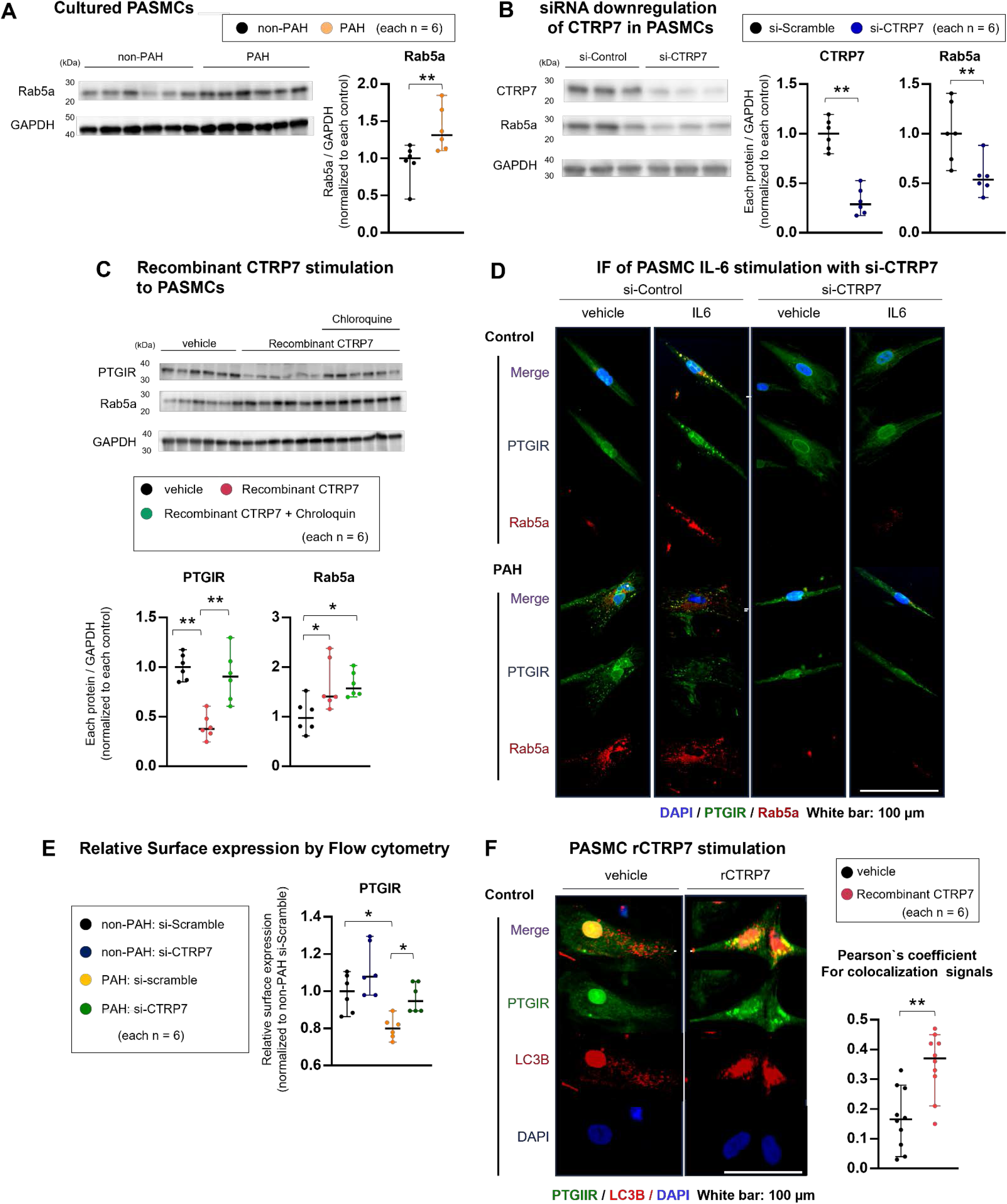
CTRP7 Downregulates Prostacyclin Analog Receptor via Rab5a in PASMCs. **(A)** Western blot and quantification of Rab5a in PASMCs from patients with PAH and from non-PAH individuals (n = 6 each). **(B)** Western blot and quantification of CTRP7, Rab5a, and GAPDH in PASMCs transfected with si-CTRP7 or si-Scramble (n = 6 each). **(C)** Analysis of representative western blot and quantification of CTRP7, Rab5a, and GAPDH in PASMCs treated with recombinant CTRP7 (1000 μM), lysosome inhibitor chloroquine (100 μM), or vehicle (n = 6 each). **(D)** Representative immunofluorescent images of PTGIR (green), Rab5a (red), and DAPI (blue) in PASMCs treated with IL-6 or vehicle, along with transfection of si-CTRP7 or si-Scramble for 24 h. Scale bar = 100 μm. **(E)** Flow cytometry analyses and quantification of cell surface expression of PTGIR in PASMCs transfected with si-CTRP7 or si-Scramble (n = 6 each). **(F)** Immunofluorescent images of PTGIR (green), LC3B (red), or DAPI (blue) in PASMCs treated with recombinant CTRP7 (1000 μM) or vehicle (n = 6 each) for 24 hours. Pearson’s correlation coefficient between PTGIR and LC3B signals. PASMCs, pulmonary artery smooth muscle cells; PAH, pulmonary arterial hypertension; CTRP7, C1q/TNF-related protein 7; PTGIR, prostacyclin analog receptor; IL-6, Interleukin-6; Rab, Ras analog in brain; LC3, microtubule-associated protein 1A/1B-light chain 3. Data are presented as mean ± SEM. Statistical analyses were performed using the 2-tailed Student’s t-test, Welch’s t-test, 1-way or 2-way ANOVA, followed by Tukey’s honestly significant difference test for multiple comparisons. Significance levels are indicated as *p < 0.05, **p < 0.01, and ***p < 0.001.

Immunofluorescence analyses showed that both PAH and non-PAH PASMCs incubated with IL-6 exhibited elevated Rab5a expression compared with that in non-PAH PASMCs (**Figure 5D**). Importantly, clear PTGIR staining was observed around the cell surface in non-PAH PASMCs. In contrast, images of PAH or non-PAH PASMCs incubated with IL-6 suggested intracellular accumulation of PTGIR and Rab5a, which was attenuated by siRNA-mediated downregulation of CTRP7 (**Figure 5D**). To quantitatively assess the cell surface expression of PTGIR, an indicator of the internalization rate, we conducted flow cytometry with cultured PASMCs, which revealed that CTRP7 downregulation increased cell surface PTGIR expression in PAH PASMCs, indicating a reduction in internalization (**Figure 5E**).

Following internalization, PTGIR is directed to the lysosomal degradation process.^24^ In line with this, co-incubation with chloroquine (CLQ) (100 μM), a lysosome inhibitor, counteracted the reduction in PTGIR expression induced by recombinant CTRP7 co-incubation (**Figure 5C**). Furthermore, immunofluorescence analyses demonstrated that the colocalization of PTGIR and LC3B, a marker of lysosomal degradation, was significantly increased by recombinant CTRP7 stimulation, suggesting that CTRP7 upregulation enhanced the lysosomal degradation of PTGIR **(Figure 5F**).

### CTRP7 Regulates the Responsiveness to MRE 269, an Active Form of Selexipag

Selexipag, an oral prostacyclin receptor agonist and a potent vasodilator, is approved for the treatment of patients with PAH or Chronic thromboembolic pulmonary hypertension.^4^ Selexipag is rapidly absorbed from the gastrointestinal tract and metabolized to its active form, corresponding to MRE269, which was used in our basic experiments.^25^ We assessed the responsiveness to MRE269 by measuring phospho–regulatory subunit 2 of protein kinase A (PKAR2) phosphorylation, as described previously.^30^ MRE269-induced PKAR2 phosphorylation was significantly higher in non-PAH than in PAH PASMCs (**Figure 6A**). Importantly, downregulation of CTRP7 exacerbated the MRE269-induced PKAR2 phosphorylation in both PAH and non-PAH PASMCs (**Figure 6A**).

**Figure 6.**
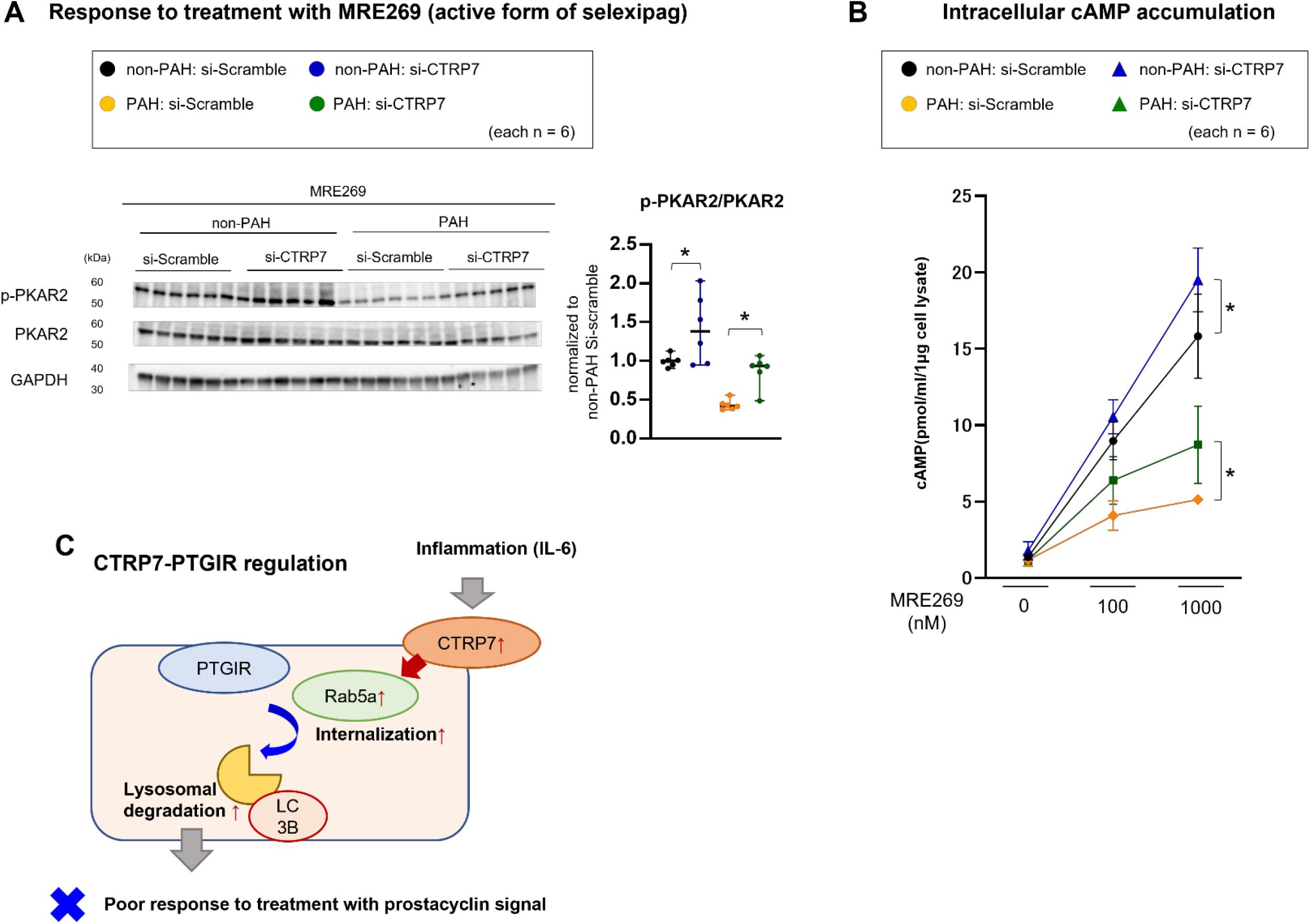
CTRP7 Regulates Responsiveness to Selexipag. **(A)** Representative western blot and quantification of p-PKAR2, PKAR2, and GAPDH in PASMCs treated with MRE269 (1000 nM) or vehicle for 24 h in patients with PAH or in non-PAH individuals, along with transfection with si-CTRP7 or si-Scramble. **(B)** Intracellular cAMP accumulation in PASMCs from patients with PAH or from non-PAH individuals treated with MRE269 (0, 100, or 1000 nM for 24 hours) and transfected with si-CTRP7 or si-Scramble. **(C)** Schematic diagram of the proposed mechanism. CTRP7 is upregulated by IL-6 and promotes the internalization and lysosomal degradation of PTGIR by regulating Rab5a expression, leading to a reduced responsiveness to PGI2 analogs. PASMCs, pulmonary artery smooth muscle cells; PAH, pulmonary arterial hypertension; CTRP7, C1q/TNF-related protein 7; PTGIR, prostacyclin analog receptor; IL-6, interleukin-6; PGI2, prostaglandin I2; cAMP, cyclic adenosine monophosphate; MRE 269, 4-[(5,6-Diphenylpyrazin-2-yl)(propan-2-yl)amino]butoxy}acetic acid Results are expressed as mean ± SEM. Statistical analyses were performed using 2-way ANOVA followed by Tukey’s honestly significant difference test for multiple comparisons. Significance levels are indicated as *p < 0.05, **p < 0.01, and ***p < 0.001.

To further assess the effect of selexipag, we measured intracellular cAMP accumulation in cultured PASMCs incubated with MRE269. Consistent with the results for PKAR2 phosphorylation, MRE269-induced intracellular cAMP accumulation was lower in PAH PASMCs compared with that in non-PAH PASMCs, irrespective of the concentration (100 nM or 1000 nM) (**Figure 6B**). Furthermore, CTRP7 downregulation significantly enhanced cAMP accumulation in both PAH and non-PAH PASMCs (**Figure 6B**). These data indicate that CTRP7 plays a role in regulating the responsiveness to selexipag via the internalization and degradation of PTGIR (**Figure 6C**).

### CTRP7 Downregulates the Response to Prostaglandin I2 Analogs in a PH Rodent Model

To investigate whether the attenuation of CTRP7 could restore PTGIR expression and improve responsiveness to selexipag in vivo, we designed a recombinant adeno-associated virus (AAV) carrying EGFP under the cytomegalovirus promoter (AAV6-shCTRP7) to downregulate CTRP7 expression in the pulmonary arteries (**Supplemental Figure IVA**), as previously described.^26^ AAV6-shCTRP7 was administered to hypoxic mice via intratracheal instillation 4 weeks before lung harvest, following which lung-specific downregulation of CTRP7 was analyzed (**Figure 7A**). This approach led to a decrease in CTRP7 expression in both the lung homogenates and plasma harvested from the mice (**Figure 7B**). Immunofluorescent images revealed that PTGIR expression in the pulmonary artery smooth muscle layer was reduced in hypoxic mice compared to that in normoxic mice, whereas PTGIR expression was increased (**Figure 7C**). Furthermore, Rab5a expression was significantly elevated in the pulmonary artery smooth muscle layer of hypoxic mice, and was attenuated by AAV6-shCTRP7 transduction (**Figure 7C**). Elastica Masson (EM) staining showed that AAV6-shCTRP7 transduction did not alter pulmonary arterial remodeling in hypoxic mice (**Supplemental Figure VB**).

**Figure 7.**
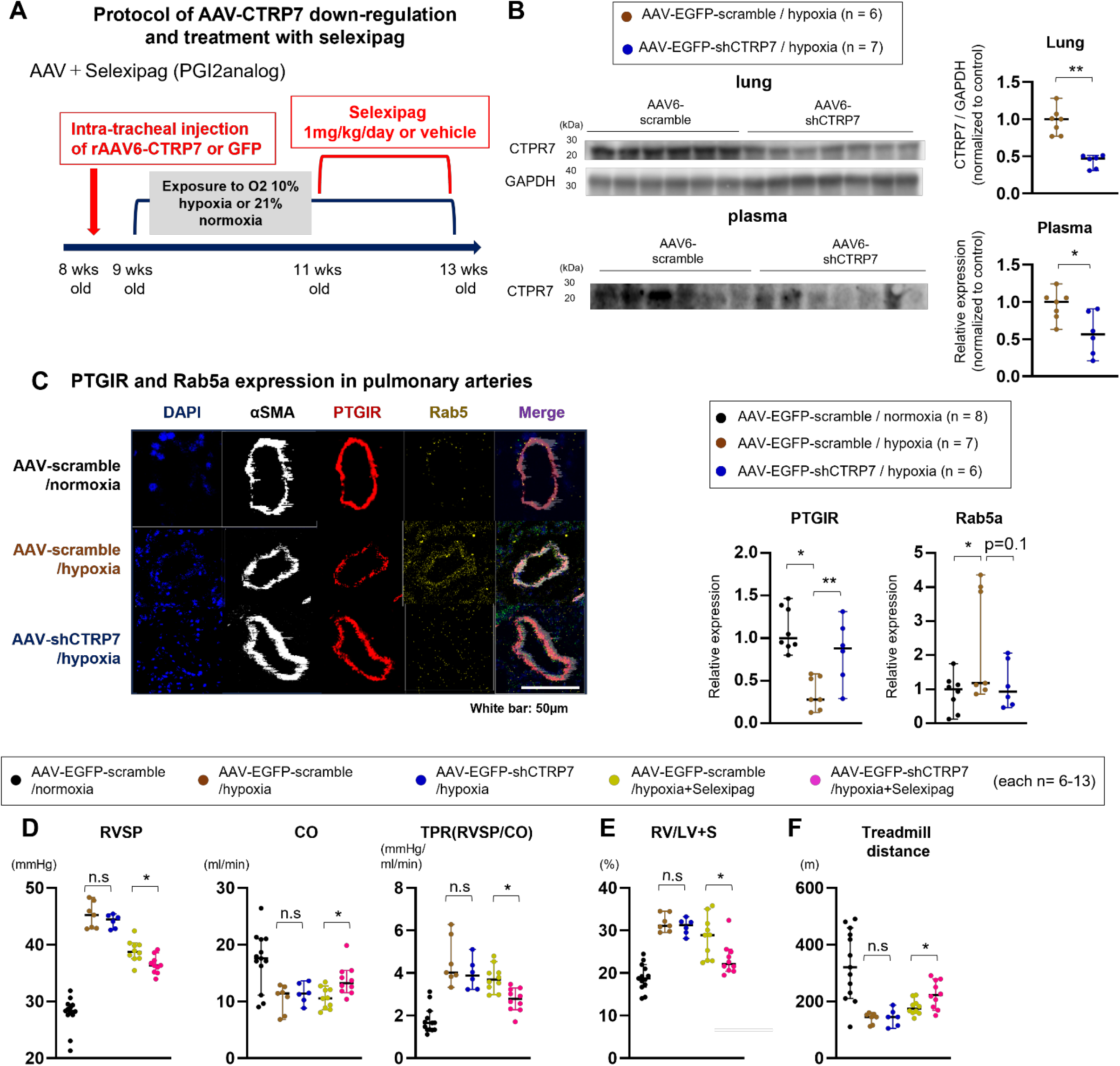
Down regulation of CTRP7 Improved the Response to Treatment with Selexipag in Hypoxic Mice. **(A)** Experimental timeline for AAV6-shCTRP7 and treatment with selexipag (1 mg/kg/day gavage) in a hypoxia PH mouse model. **(B)** Western blot and quantification of CTRP7 in the lungs or plasma harvested from hypoxic mice treated with AAV6-shCTRP7 or AAV6-scramble via intratracheal instillation (n = 6–7). **(C)** Representative immunofluorescent images of αSMA (white), PTGIR (red), Rab5a (yellow), and DAPI (blue) of the distal PAs in hypoxic mice (10% O_2_ for 4 weeks) with downregulation of CTRP7 using AAV6-shCTRP7. Scale bar = 50 μm. **(D)** RVSP was assessed using a catheter. Ultrasonography was performed to assess CO. TPR was calculated as RVSP per CO. (**E**) Right ventricular hypertrophy was assessed by the weight ratio of the right ventricle to the left ventricle plus the septum. (**F**) Exercise capacity was assessed using the workload of the treadmill exercise. AAV6, adeno-associated virus 6; PH, pulmonary hypertension; CTRP7, C1q/TNF-related protein 7; PTGIR, prostacyclin analog receptor; PAs, pulmonary arteries; RVSP, right ventricular systolic pressure; CO, cardiac output; TPR, total pulmonary resistance; αSMA, alpha smooth muscle actin. Results are expressed as mean ± SEM. Comparisons of parameters were performed using the 2-tailed Student t-test, Welch t-test, 1-way or 2-way ANOVA, followed by the Tukey’s honestly significant difference test for multiple comparisons. *p < 0.05. **p < 0.01. ***p < 0.001.

In terms of hemodynamics of mice, transduction using AAV6-shCTRP7 did not alter PH, heart rate, or blood pressure (**Figure 7D** and **Supplemental Figure IVC**). This was evidenced by increased right ventricular systolic pressure (RVSP), total pulmonary artery resistance (RVSP/cardiac output [CO]) (**Figure 7D**), right ventricular hypertrophy (weight ratio of the right and left ventricle (RV/LV) + septum) (**Figure 7E**), LV and RV dilation or function (right ventricular diastolic dimension (RVDd), tricuspid annular plane systolic excursion (TAPSE), right ventricular fractional area change (RVFAC), and pulmonary acceleration time per ejection time (PAT/ET) (**Supplemental Figure VD** and **VE**), and lowered exercise capacity (**Figure 7F**).

To evaluate the role of CTRP7 in responsiveness to selexipag, hypoxic mice were administered AAV6-shCTRP7 at three-week intervals, followed by treatment with selexipag (1 mg/kg/day) via gavage (**Figure 7A**). Selexipag improved RVSP without significant CTRP7 alteration by AAV transduction, whereas the CO, total pulmonary resistance (TPR), RV hypertrophy, and exercise capacity remained comparable (**Figure 7D-F**). Transduction of AAV6-shCTRP7 significantly enhanced selexipag-induced improvement in RVSP in hypoxic mice (**Figure 7D**). Similarly, hypoxic mice treated with AAV6-shCTRP7 and selexipag exhibited additional improvements in the CO, TPR, RV hypertrophy, and exercise capacity (**Figure 7D-F**). These results suggest that CTRP7 plays a role in diminishing the responsiveness to selexipag without significantly impacting the progression of pulmonary arterial remodeling.

## Discussion

The key findings of this study after combining data from both human and basic analysis were as follows:1) CTRP7 exhibited elevated levels in PASMCs and plasma of patients with PAH; 2) inflammation induced an increase in CTRP7 and a decrease in PTGIR via the upregulation of Rab5a-mediated internalization and lysosomal degradation; and 3) higher CTRP7 levels were associated with a reduced response to PGI2 analogs in a mouse model of PH.

### Higher CTRP7 Expression in PASMCs Correlates with Elevated Plasma Levels in Patients with PAH

CTRP7, a member of the CTRP cytokine family that is primarily associated with glucose metabolism and inflammation, is predominantly expressed in the lungs.^14^ However, its role in the pulmonary vascular function remains unclear. This study revealed the intricate molecular mechanisms that connect inflammation to pulmonary vasoreactivity via the CTRP7-PTGIR axis.

The primary objective of this study was to identify candidate biomarkers capable of predicting patient responsiveness to pulmonary vasodilators. The two-step approach used in the present study involved RNA sequencing of PASMCs and assessment of plasma levels. Notably, CTRP7 emerged as a prominent biomarker, displaying increased expression in both PASMCs and plasma obtained from patients with PAH (**Figure 1A-C**). Hypoxic mice exhibited elevated CTRP7 levels in their lungs and plasma, but not in their liver or adipose tissue (**Supplemental Figure IB**). Moreover, the AAV-mediated downregulation of CTRP7 in the lungs of mice was linked to reduced plasma CTRP7 levels (**Figure 7B**). These findings suggest that plasma CTRP7 levels may serve as a reliable indicator of CTRP7 expression in the pulmonary arteries of patients.

Challenges persist in the quest to predict protein expression in the pulmonary arteries using plasma samples, owing to the complex nature of protein secretion in the vasculature and other organs, as well as the heterogeneous pathogenesis of PAH. Considering that lung or pulmonary artery biopsies, categorized as class III in the ESC/ERS guidelines,^4^ pose potential lethal risks to PAH patients, the investigation of target factors using tissue and plasma samples, as demonstrated in this study, is imperative. Importantly, additional markers or tests, such as IL-6 or other inflammatory markers, could enhance the precision and effectiveness of estimating CTRP7 expression in the pulmonary arteries based on plasma levels (**Figure 3A and 3B**).

### Inflammation Triggers an Increase in CTRP7 Expression and a Concurrent Decrease in Prostacyclin Analog Receptor via Upregulation of Rab5a-mediated Internalization and Lysosomal Degradation

Inflammation plays a pivotal role in the development of PAH, with factors such as monocyte chemoattractant protein-1 (MCP1), tumor necrosis factor alpha (TNFα), IL-1β, and IL-6 acting as key mediators.^31, 32^ These factors influence the responsiveness to vasodilators; studies in animal models have shown that immunosuppressive therapies and IL-6 blockade (e.g., tocilizumab) can ameliorate PH.^33^ A similar clinical benefit was observed, particularly in patients with CTD-PAH, although the underlying mechanisms remained obscure.^34^

We conducted ChIP analyses and RNA sequencing, which provided clear evidence that IL-6 upregulates CTRP7 expression in PASMCs via transcriptional regulation (**Figure 3D**).

Subsequently, CTRP7 downregulated the expression of PTGIR by promoting its internalization via a Rab5-mediated mechanism, ultimately resulting in a poor response to selexipag both in vitro and in vivo. These patterns were consistently observed in various PAH models, such as hypoxic mice and rats, and in monocrotaline-or Sugen/hypoxia-induced PH models (**Figure 2D**). PTGIR is a G-protein-coupled receptor; PGI2 selectively binds to and activates PTGIR to carry out its physiological actions.^35^ The expression of PTGIR is influenced by various factors, including transcriptional regulation, agonist-induced desensitization, internalization, or degradation.^35^ While previous studies have demonstrated the internalization of PTGIR by Rab5, its mechanisms in the pulmonary arteries of patients with PAH were unclear. As mentioned above, this study is the first to demonstrate that inflammation promotes Rab5-mediated internalization and degradation of PTGIR via CTRP7 in the PASMCs of patients with PAH.

### Higher CTRP7 Levels are Associated with a Reduced Response to Prostaglandin I2 Analogs in a Mouse Model of Pulmonary Hypertension

To assess the role of CTRP7 in the pulmonary arteries in vivo, we employed AAV-mediated CTRP7 alterations in hypoxic mice. Treatment with selexipag partially ameliorated PH in the hypoxic PH model mice, resulting in a relative decrease in RVSP, while no significant changes were observed in the CO, TPR, or exercise capacities (**Figure 7D-F**). AAV6-mediated downregulation of CTRP7 in the pulmonary arteries significantly enhanced selexipag-induced improvement of PH in hypoxic mice (**Figure 7D**), consistent with the results of in vitro experiments using cultured PASMCs (**Figure 6**). These findings indicate that altering CTRP7 expression can enhance responsiveness to pulmonary vasodilators, irrespective of other conditions such as inflammation or sex.

In a clinical setting, therapies aimed at suppressing IL-6, a key mediator of inflammation, have demonstrated benefits in various diseases, including rheumatoid arthritis^33^ and scleroderma.^36^ However, for patients with PAH, therapeutic blockade of IL-6 (e.g., tocilizumab) did not lead to improvements in pulmonary vascular resistance (PVR).^37^ These results suggest that it may be worthwhile to shift our focus toward CTRP7, acknowledging its role beyond inflammation in the context of treatments involving pulmonary vasodilators. Although further research is required to develop novel treatment strategies that combine anti-inflammatory and vasodilatory approaches, CTRP7 has emerged as a central focus in the therapeutic management of PAH.

### Clinical Implication: Plasma Assessment of CTRP7 for Monitoring Pulmonary Vasoreactivity

Compared to the accumulation of basic research on pulmonary artery remodeling, the mechanisms underlying pulmonary arterial responsiveness to vasodilators in clinical settings require further investigation. Notably, the prostacyclin signaling pathway remains poorly understood compared to the well-established NO-sGC-cGMP axis. Previous studies have focused on sex differences considering the higher prevalence of PH in female patients. For instance, estrogen is known to enhance the expression of vasodilatory prostacyclin receptors in the vasculature,^26^ and male patients treated with epoprostenol showed poorer outcomes than their female counterparts.^27^ From another perspective, sub-analyses of the phase 3 GRIPHON study, which assessed the efficacy and safety of selexipag in patients with PAH, have suggested that the influence of cardiovascular comorbidities on the treatment effect of selexipag is relatively small or unclear.^40^ Ultimately, existing basic research has not provided sufficient evidence for the molecular mechanisms of pulmonary arterial responsiveness to prostacyclin analogs.

Similarly, in clinical practice, the current guidelines for selecting pulmonary vasodilators for the management of PAH do not incorporate molecular evidence, and may overlook the variable phases and conditions that patients may experience during the clinical course of the disease. Pulmonary vasoreactivity demonstrates that a patient’s responsiveness to treatment can change over time or with age. Existing vasoreactivity tests, such as NO or iloprost inhalation and intravenous adenosine, may indicate a potential response to calcium channel blocker therapy but fall short in reliably predicting responses to other vasodilators.

The results of the present study suggest that plasma CTRP7 levels may serve as a valuable tool for monitoring various phases of PH. In the in vivo experiments, levels of CTRP7 significantly increased in both plasma and lung tissue during the early stages of hypoxic exposure or following monocrotaline stimulation, which closely correlated with elevated IL-6 levels (**Figure 4B-C** and **Supplemental Figure IVA**). This pattern may resemble the onset of PAH or the active phase of connective tissue diseases in clinical settings. Subsequently, these reactions decreased, but remained elevated during the chronic phase of hypoxic conditions. These findings underscore the potential of assessing plasma CTRP7 levels, either independently or alongside other inflammatory indicators, to predict pulmonary arterial expression and responsiveness to selexipag in clinical practice.

### Limitations

This study has several limitations. First, we did not investigate the role of CTRP7 in pulmonary artery endothelial cells. Secondly, AAV transduction was not specifically targeted to the smooth muscle layer in the pulmonary arteries. Additionally, the study did not assess the role of CTRP7 in different animal models, such as monocrotaline-or SU5416/hypoxia-induced PH model rats.

### Conclusions

In conclusion, in this study, we identified CTRP7 as a potential predictor of responsiveness to PGI2 analogs based on findings from experiments involving cells, animal models, and the plasma of patients with PAH. The results suggest that CTRP7 may serve as a novel biomarker for predicting the response to PGI2 analogs.

## Acknowledgments

We are grateful to the lab members in the Department of Cardiovascular Medicine at Tohoku University for valuable technical assistance, especially Yumi Watanabe, Hiromi Yamashita, and Kaori Miyamura.

